# Defective ORF8 dimerization in delta variant of SARS CoV2 leads to abrogation of ORF8 MHC-I interaction and overcome suppression of adaptive immune response

**DOI:** 10.1101/2021.08.24.457457

**Authors:** Armi M Chaudhari, Indra Singh, Madhvi Joshi, Amrutlal Patel, Chaitanya Joshi

**Author notes:** **Equal Contribution:** Authors had contributed equally (Armi M Chaudhari, Indra Singh). **Corresponding author:** Chaitanya Joshi, Gujarat Biotechnology Research Centre (GBRC), Department of Science and Technology, Government of Gujarat, 6th Floor, Block B&D, MS Building, Gandhinagar-382011.; **E-mail:**.

## Abstract

In India, the breakthrough infections during second wave of COVID-19 pandemic was due to SARS-COV-2 delta variant (B.1.617.2). It was reported that majority of the infections were caused by the delta variant and only 9.8% percent cases required hospitalization whereas, only 0.4% fatality was observed. Sudden dropdown in COVID-19 infections was observed within a short timeframe, suggesting better host adaptation with evolved delta variant. Down regulation of host immune response against SARS-CoV-2 by ORF8 induced MHC-I degradation has been reported earlier. The Delta variant carried mutations (deletion) at Asp119 and Phe120 amino acids which are critical for ORF8 dimerization. The deletions of amino acids Asp119 and Phe120 in ORF8 of delta variant results in structural instability of ORF8 dimer caused by disruption of hydrogen bonding and salt bridges as revealed by structural analysis and MD simulation studies of ORF8 dimer. Further, flexible docking of wild type and mutant ORF8 dimer revealed reduced interaction of mutant ORF8 dimer with MHC-I as compared to wild type ORF8 dimer with MHC-1, thus implicating its possible role in MHC-I expression and host immune response against SARS-CoV-2. We thus propose that mutant ORF8 may not hindering the MHC-I expression thereby resulting in better immune response against SARS-CoV-2 delta variant, which partly explains the sudden drop of SARS-CoV-2 infection rate in the second wave of SARS-CoV-2 predominated by delta variant in India

**Graphical Abstract:** 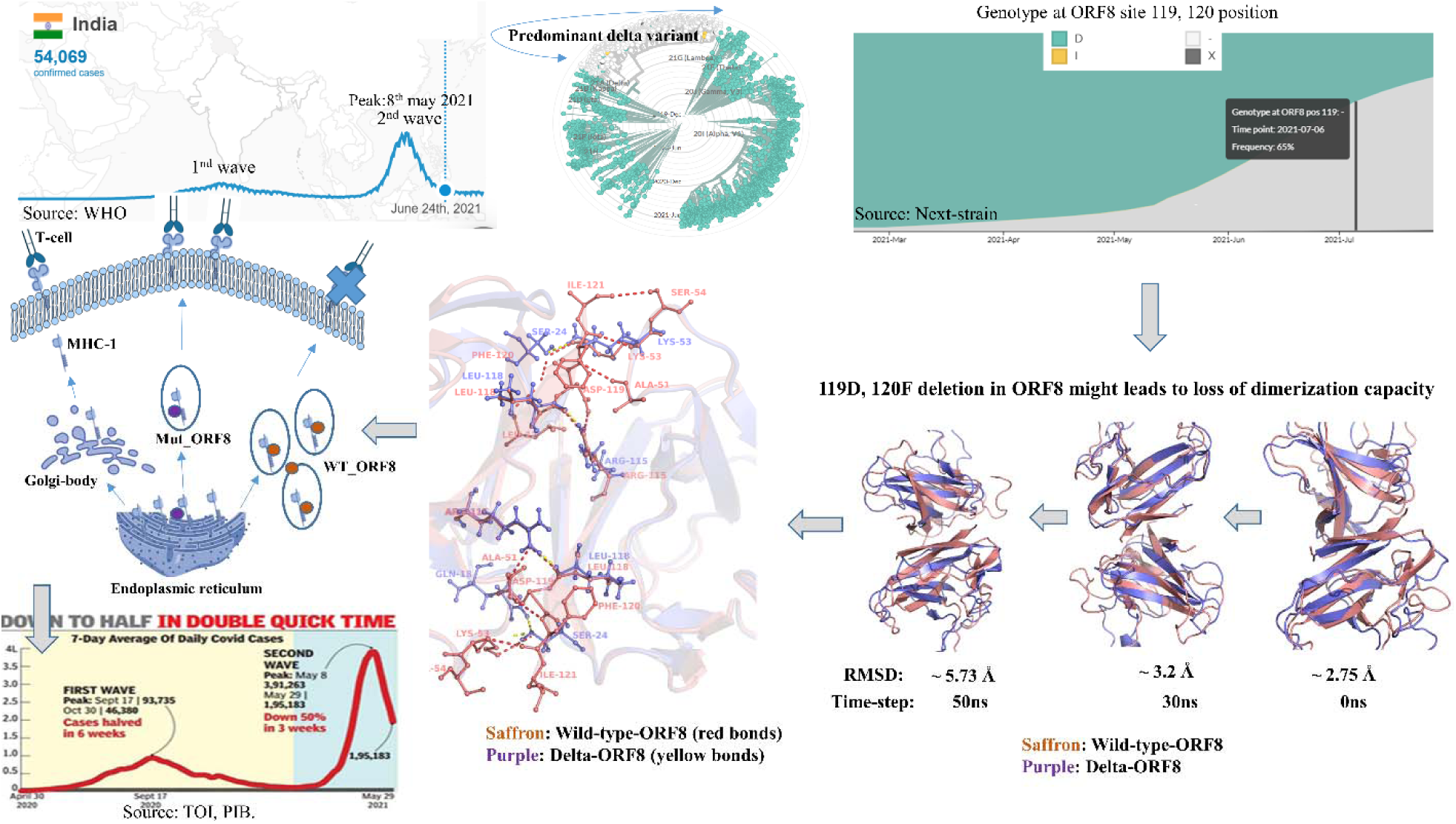

## 1 Introduction

SARS-CoV-2 pandemic had infected more than 199 million people and more than 4 million deaths worldwide till 4^th^ August 2021. During this pandemic, virus had mutated to evade the host immune system and also to enhance its transmission. These variants were detected using high throughput sequencing methods and their effect on virus is studied extensively. With these evolving variants, SARS-CoV-2 Interagency Group (SIG) of US government come up with Variant Classification scheme that defines three classes of SARS-CoV-2 variants, such as 1) VOI 2) VOC and 3) VOHC. Among them, delta variant belonging to the group of VOCs had surged to sudden increases in infection during second wave in India. This delta variant is seeming to be highly contagious due to mutations in spike. Several other mutations like D614G in modulating higher spike infectivity and density, E484K for decreased antibody neutralization, N501Y and K417N for altering spike interacting with ACE receptor and antibodies derived from human were reported. [1–3]. Recent reports suggests that NTD (N-Terminal Domain) is known to be supersite for antibody mediated binding[4–6]. Reports on rigidization in NTD of spike had led to the antibody escape mechanism in this delta variant [7]. These examples are enough to show case how lethal this variant is in terms of transmittance, infectivity and evading host immune responses. Opposite to the same, some rare mutations like C241T was favoring host also [8]

In India second wave was persisted from middle of the march 2021, till June 2021[9]. Preliminary focus of this research lies on finding possible reason of sudden drop down of second wave of SARS-CoV-2 in halved period compared to first wave with increased seroprevalence. Virus genome is extensively studied and possible mutations favoring host were identified using protein dynamics approach, among them ORF8 carrying mutations Δ119Asp and Δ120Phe had grabbed our attention due to their direct involvement in dimerization of ORF8 by forming hydrogen bonds and Salt-bridges. Crystal structure of ORF8 reported was taken as a reference structure for analyzing effect of these deletions using molecular modelling and simulation approach [10]. ORF8 is known to be important protein for SARS-CoV-2 mediated infection by down regulation MHC-I molecule in ER (endoplasmic reticulum pathway) mediated protein trafficking pathway [11]. ORF8 involvement in endoplasmic reticulum mediated stress and antagonizing IF-beta (interferon beta) for immune evasion is also known [12]. Deletion of ORF8 leads to decreased severity of infection as reported [13,14]. These examples show case the involvement of ORF8 in modulating host immune response and majorly by downregulating MHC-I. Exact interface of MHC-I binding with ORF8 is not known yet. In this study effect of these Δ119Asp and Δ120Phe deletions with respect to ORF8 dimerization is studied. Flexible induced docking was performed to study the ORF8 mediated MHC-I binding. Mutations were correlated with timeline of second wave and available cohort study on seroprevalence.

## 2 Material and Methods

### 2.1 Data retrieval

Crystal structure of ORF8 (PDB ID: 7JTL) protein of SARS-CoV-2 (WT_ORF8) was retrieved from protein data bank [10]. Protein sequence ORF8_GBRC_NCD_370 of SARS-CoV-2 delta variant (MUT_ORF8) was obtained from inhouse sequencing (Sequence submitted to GAISAD with accession number EPI_ISL_2001211) and fasta sequence of MHC-I protein (Accession no: NP_005505.2) was downloaded from NCBI.

### 2.2 Protein structure modelling and Molecular Dynamics Simulations studies

3-Dimentional structures of MUT_ORF8 protein as well as of MHC-I protein were built using homology modelling panel under the Schrodinger suite release 2021-2 [15]. The fasta sequences of MUT_ORF8 and MHC-I protein were imported into the Schrodinger suite. Homology blast search resulted in the templates 7JTL and 6AT5 corresponding to MUT_ORF8 and MHC-I respectively. Protein preparation wizard was then used for the refinement of protein structures. Additionally PRIME module was also used to add missing residues and pKa refinement of proteins was done using epic module of Schrodinger suite [16].

Conformational stability of WT_ORF8 and MUT_ORF8 dimers were inspected using molecular dynamic simulations studies in detail using DESMOND module implemented in Schrodinger suite 2021-1 till 200 nanoseconds (ns)[17]. OPLS4 force field was applied to refine the WT_ORF8 and MUT_ORF8 dimeric proteins as well as H-bonds were refilled using structure refinement panel implemented in Schrodinger suite [18,19]. Particle mesh Ewald method was applied for calculation of long-range electrostatic interactions [20]. Also, at every 1.2 ps intervals the trajectories were recorded for the analysis. The proteins WT_ORF8 and MUT_ORF8 were placed in the center of the dodecahedron water box of the TIP3P water model of size wild 353968Å and 360038Å respectively [21]. The whole system was neutralized using 1.5 mM Salt concentration. A coupling constant of 2.0 ps under the Martyna–Tuckerman–Klein chain-coupling scheme was used for pressure control and the Nosé–Hoover chain-coupling scheme at 310.3K was used for temperature control of the system [22]. The whole system was initially energy minimized by steepest descent minimization. Total negative charges on the protein structures of WT_ORF8 and MUT_ORF8 were balanced by appropriate number of Na+ ions to make the whole system neutral. Further, energy-minimized protein structures were subjected to position restrained dynamics for 200 ns, allowing water molecules to equilibrate and the whole protein system was kept fixed. Optimized system was subjected to MD run for 200 ns at 310.5 K and 1 atmospheric pressure (NPT ensemble). The binding energy of the system was calculated for each of the protein structures and stability of complex was monitored by analyzing RMSD, RMSF, radius of gyration and H-bonds of each dimer throughout simulation run time. High resolution images were generated using Pymol and biovia Discovery studio (BIOVIA, Dassault Systèmes, BIOVIA Workbook, Release 2020; Schrodinger, LLC. 2010. The PyMOL Molecular Graphics System). Protein networking was studied into NASP server available online [25]. Ramachandran plots were generated into zlab Ramachandran plot server [26].

### 2.3 Binding energy (MMGBSA) Calculation

The binding free energy of WT_ORF8 and MUT_ORF8 dimers were calculated by Prime Molecular Mechanics-Generalized Born Surface Area (MMGBSA) using thermal_mmgbsa.py implemented under PRIME module of Schrodinger suite [27–29]. The binding free energy of each protein provides a summary of the biomolecular interactions between monomeric chains of protein dimer. OPLS4 force-field and VSEB solvation model were used for MMGBSA calculation. The binding energy includes potential energy as well as polar and non-polar solvation energies were calculated as following.

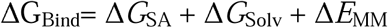

### 2.4 Principal Component analysis (PCA) and Dynamics cross-correlation matrix (DCCM) calculation

To perform PCA, Primarily the covariance matrix C was calculated. The eigenvectors and eigenvalues were obtained for the covariance matrix C [30]. The principal components (PCs) are projections of a trajectory on the principal modes, of which usually the first few ones are largely responsible for the most important motions. The elements Cij in the matrix C are defined as:

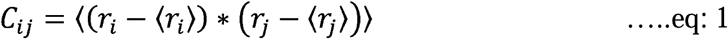

From equation 1, ri and rj are the instant coordinates of the ith or jth atom, ⟨*r*_*i*_⟩ and ⟨*r*_*j*_⟩ and mean the average coordinate of the ith or jth atom over the ensemble.

Correlative and anti-correlative motions are playing a key role in the recognition as well as binding in the biological-complex system. These motions can be prevailed through molecular dynamics simulation trajectories by defining the covariance matrix about atomic fluctuation. The magnitude of correlative motions of two residues can be represented by the cross-correlation coefficient, Cij. It is defined by following equation:

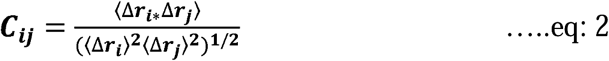

Here, i (j) is ith (jth) two residues (or two atoms/proteins), Δri (Δrj) is the displacement vector corresponding to ith (jth) two residues (or two atoms/proteins), and ⟨.. ⟩ is for the ensemble average. The value of Cij ranges from +1 to -1. +Cij denotes positive correlation movement (same direction) shown in blue color, and -Cij denotes anti-correlation movement (opposite direction) shown in red color. The higher the absolute value of Cij is, the more correlated (or anti-correlated) the two residues (or two atoms or proteins). PCA and DCCM both were evaluated by using run *trj_essential_dynamics*.*py*, a python script under Desmond module of Schrodinger 2021-1[31].

## 3 Results

### 3.1 Effect of deletions on the binding affinity of MUT_ORF8 dimer

WT_ORF8 protein comprises of two monomeric chains existing in the form of a dimeric structure which is tightly packed with the help of various electrostatic interactions and H-bonds (Supplementary figure S1). The key residues involved in the packing of WT_ORF8 dimers are Lys53, Arg115, Asp119, Phe120 and Ile121. Other residues involved in intra chain bonds between dimers of WT_ORF8 are Gln18, Ser24, Ala51, Arg52 and Ser54 (Figure 1A). These dimers are closely held together with four salt bridges formed between A: Asp119-B: Arg115, A: Arg115-B: Glu92, B: Asp119-A: Arg115 and B: Arg115-A: Glu92. Other interactions are several H-bonds between Phe120 and Lys53, Lys53 and Ser24, Gln18-Lue22, Arg52 and Ile121 (Figure 1A). In WT_ORF8, amino acids Asp119 and Phe120 are predominantly involved in the formation of salt bridges as well as Hydrogen bonds (Supplementary figure S1: C & D). The detailed analysis of MUT_ORF8 dimer protein showed that its monomer is attached with each other with only 1 salt bridge between C: Arg15-D: Glu92. Six H-bonds are formed between amino acids C: Gln18-D: Ser24 (one H-bond), C: Arg115-D: Leu118 (two H-bonds) and C: Ile119-D: Ala51 (three H-bonds) (Supplementary figure S1: C & D). Protein structural network analysis shows reduced nodes (amino acids) and bonds (edges). WT_ORF have 722 edges while Mut_ORF8 have only 714 (Figure 1B). Decreased edges correlated with reduced protein-protein interactions (here in case of monomers). These decreased monomeric interactions in MUT_ORF8 might leads to less stable dimer formation of ORF8. Ramachandran plot for both variants of ORF8 is shown in figure 1C. WT_ORF8 possess majority of amino acids in highly preferred region (green) with no questionable interactions, while mutant ORF8 possess two questionable angles for amino acids C: E-64 and D: S-67 (shown in red dots), depicting decrease in protein stability of MUT_ORF8 (Figure 1C). Contact plot generated for inter and intra molecular interactions within ORF8 dimers where WT_ORF8 possess a higher inter-intra molecular interactions compared to MUT_ORF8, and leads to more stable dimer (Figure 1D). Overall structural studies of proteins suggests that WT-ORF8 seems to be stable dimer compared to MUT_ORF8 by forming strong interactions like hydrogen bonds and salt bridges, these observations were further confirmed using molecular dynamics approach.

**Figure 1:**
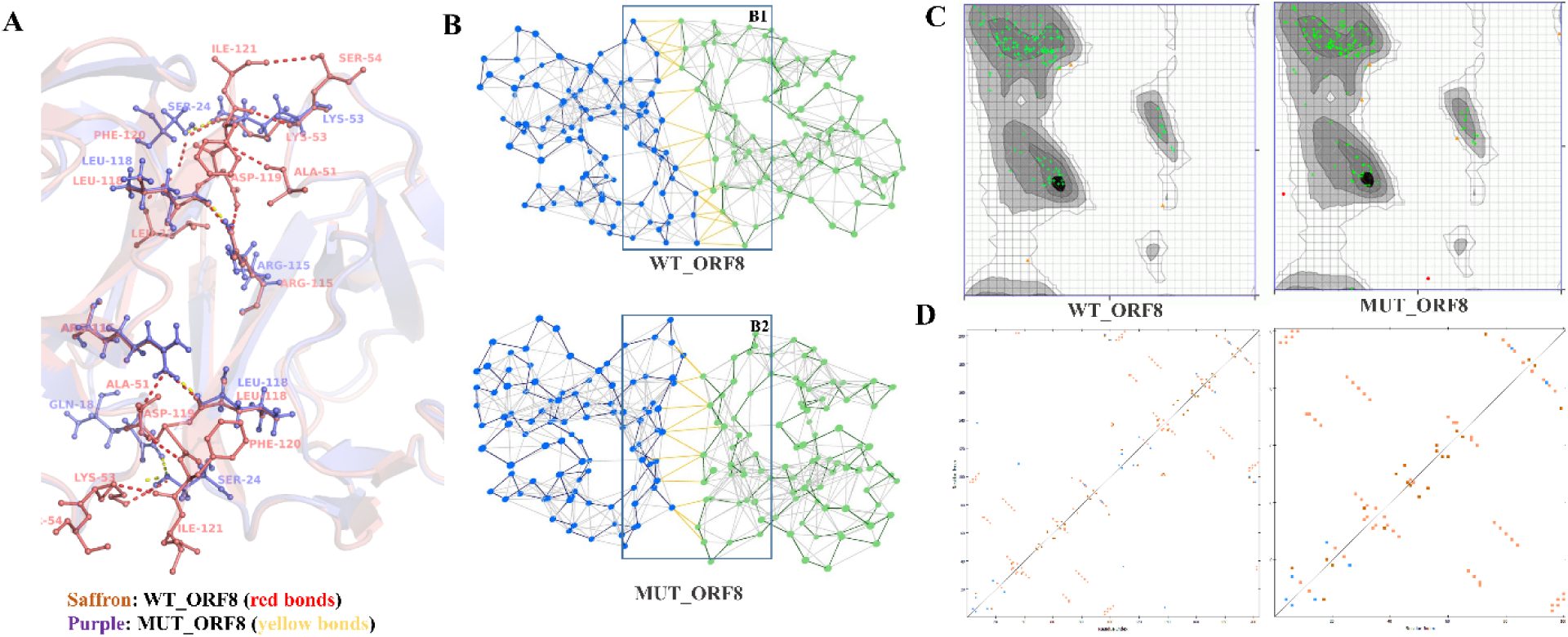
Change in bond formation within WT_ORF8 and MUT_ORF8 due to 119Asp and 120F deletion: **1A:** Superimposition of WT_ORF8 shown in saffron color and MUT_ORF8 shown in purple color. Hydrogen bond formation within two monomeric units of ORF8 is illustrated using Pymol where red and yellow bonds representing bond formation within WT_ORF8 and MUT_ORF8. **1B:** Network analysis of protein structures using NASP sever, where B1 and B2 represent network between dots as a node (amino acids) and inter and intramolecular bonds as an edge (yellow) for WT_ORF8 and MUT_ORF8 respectively. WT_ORF8 possess 203 nodes and 722 edges while MUT_ORF8 possess 202 nodes and 714 edges. **1C:** Ramachandran plot for WT_ORF8 and MUT_ORF8. Green dots represent highly preferred observations, yellow dots represent preferred observations and red dots represents questionable observations. MUT_ORF8 possess two questionable observations which are C: E-64 and D: S-67, while WT_ORF8 posses no such kind of observations. **1D:** Contact plot showing amino-acids contacts between monomeric units of WT_ORF8 and MUT_ORF8. Blue color shows main chain-side chain interactions, Saffron color shows main chain-main chain interactions, and brown color shows side chain-side chain interactions within monomeric subunits.

### 3.2 Molecular dynamics reveals breakdown/dissociation of ORF8 dimer in delta variant

After execution of the classical molecular dynamics simulations for 200ns the root mean square deviation (RMSD) of the trajectories were calculated, to identify the region of WT_ORF8 and MUT_ORF8 dimers showing deviations with respect to the initial structure. The RMSD plot clearly showed that the conformational stability of WT_ORF8 is greater than MUT_ORF8 (Figure 2C). The RMSD of MUT_ORF8 dimer is on higher side throughout the simulation run time as compared to initial conformations. The RMSD of WT_ORF8 has fluctuation between 1.527-5.652Å throughout the simulation runtime of 0-200ns. Whereas RMSD of the MUT_ORF8 is fluctuating from 1.73Å to 4.498 during 0-10ns, 5-10.47Å during 10-30ns, 10.478-12.049Å during 30-100ns and 12.049-14.79 during 100-200ns. Number of H-bonds were plotted for the duration of 0-200ns simulation time (Figure 2D), showing that WT_ORF8 has number of H-bonds between 3-22 throughout the simulation. Maximum number of bonds i.e. 22 H-bonds are formed in WT_ORF8 at 102ns simulation time. Number of H-bonds were also calculated for MUT_ORF8 varying from 0 to 15. Radius of gyration was also studied to see the compactness of protein structure of WT_ORF8 and MUT_ORF8. Δ119Asp and Δ120Phe were not favoring dimer formation in ORF8 which is seen during simulation, Supplementary video 2 shows the dissociation/breakdown of ORF8 monomers in mutant ORF8. In wild type no such breakdown occurs (See supplementary video 1) The highest radius of gyration of MUT_ORF8 throughout the simulation time, suggesting a less tight packing of MUT_ORF8 as compared to WT_ORF8 (Figure 2E). The value of radius of gyration is ranges from 18.416 -24.386 in WT_ORF8 whereas, from 18.492-25.444 in MUT_ORF8. To investigate the effect of mutation on the dynamics of the backbone atoms, RMSF values for each dimer were calculated at each time point of the trajectories. Root mean square fluctuation (RMSF) values of WT_ORF8 is shifting from 0.737 to 11.997 Å. Only Residues 67, 68, 69 and 70 of WT_ORF8 are having high RMSF value of 11.997Å, whereas other residues showing less RMSF value (Figure 2H). RMSF values for MUT_ORF8 dimer is 1.3 to 7.078Å. it is on higher side throughout simulation as compared to WT_ORF8. Dynamics cross-correlation matrix (DCCM) of WT_ORF8 and MUT_ORF8 were plotted (Figure 2F & 2G) In DCCM WT_ORF8 holding higher intensity for blue color as compared to MUT_ORF8. Positive Cij values signaling blue colors that leads to improved interaction profile between residues.

**Figure 2:**
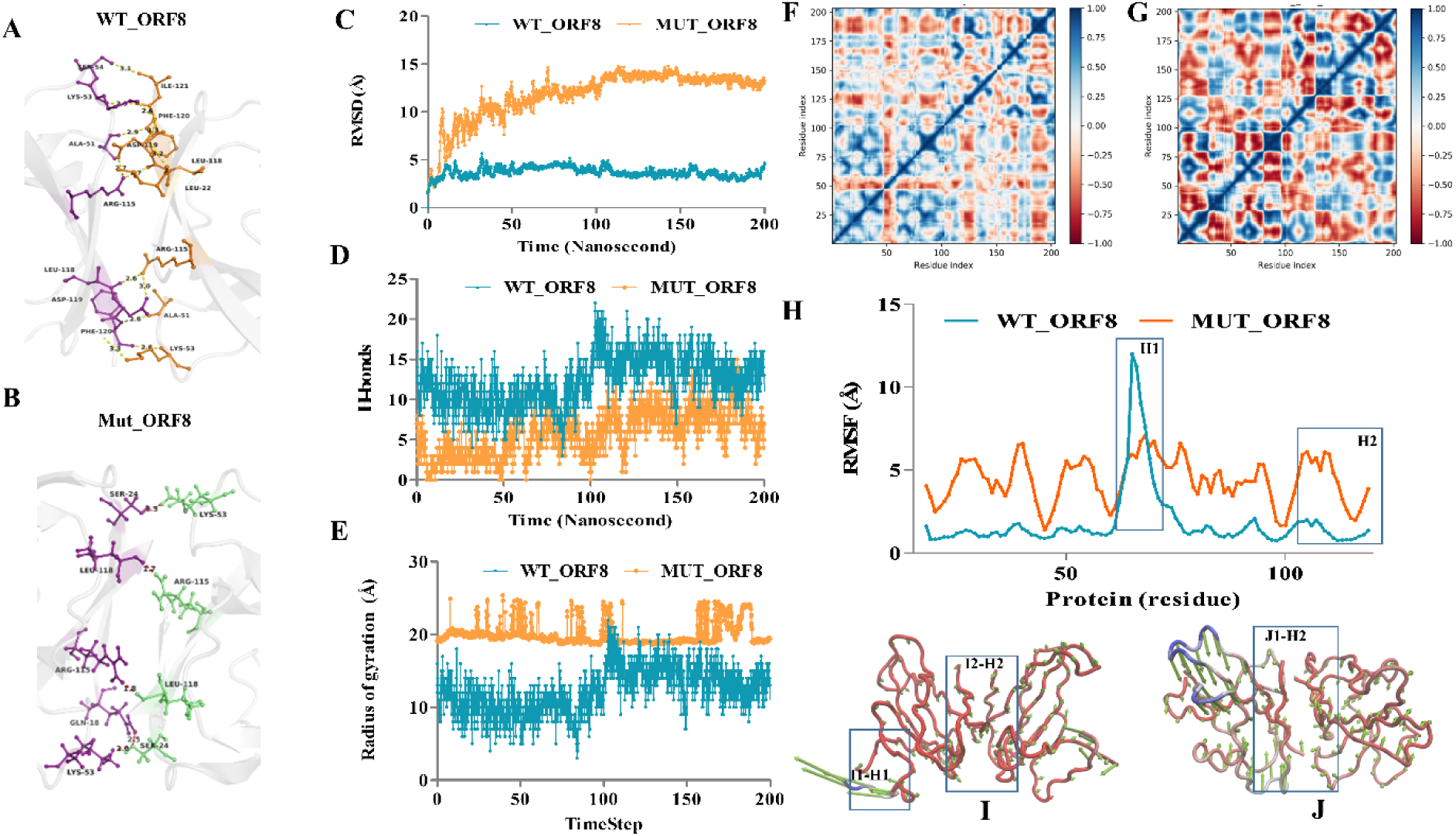
Molecular dynamics studies for both variants of ORF8 dimer. **2A:** Intramolecular interactions between WT_ORF8 monomeric subunits **2B:** Intramolecular interactions between MUT_ORF8 monomeric subunits. **2C:** RMSD (root mean square deviation) within WT-ORF8(cyan) and MUT_ORF8 (orange) complex. **2D:** Hydrogen bonds formation within WT-ORF8(cyan) and MUT_ORF8 (orange) complex. **2D:** Radius of gyration for WT-ORF8(cyan) and MUT_ORF8 (orange) complex **2F & 2G:** Dynamics cross-correlation matrix obtained from trajectories of WT_ORF8 and MUT_ORF8 complexes respectively. Blue to red color represents the cij values between 1 to -1. No cross correlation was shown by white color. **2H:** RMSF (root mean square fluctuation) in WT-ORF8(cyan) and MUT_ORF8 (orange) complex. **2I:** PCA1 mode of WT-ORF8, length of arrow is in linear relation between protein dynamics/fluctuation during trajectories blue color shows highly dynamic regions, while red color shows less dynamics regions. **2J:** PCA1 mode of MUT-ORF8.

The binding energy (MMGBSA) calculations were performed for both dimers WT_ORF8 and MUT_ORF8. From figure 3A it is clearly seen that WT_ORF8_wt is more stable having higher negative free energy as compared to MUT_ORF8. The electrostatic energy of WT_ORF8 and MUT_ORF8 was -295.08 and -97.27, respectively. Similar pattern has been observed for ΔG bind, Vander Waal energy, H-bond energy, lipophilic energy, covalent energy, and solvation energy for WT_ORF8 and MUT_ORF8 (Figure 3A). It is evident that only three amino acids i.e., Arg115, Val117 and Ile121 are involved in dimerization of MUT_ORF8 as compared to WT_ORF8 where Val114, Arg115, Val116, Val117, Lue118, Asp119, Phe120 and Ile121 are involved in the stabilization of the WT_ORF8 dimer (Figure 3C). Electrostatic potential are major energies which were contributing in dimer formation. Energies were visualized in ABPS module implemented in Pymol 1.8. As shown in figure 3B, WT_ORF8 have higher opposite attraction (positive-negative) compare to MUT_ORF8. Box B1 and B2 shows the region where these electrostatic potentials persist for both variants. Increased electrostatic potential among amino-acids of WT_ORF8 shows favorable dimer formation compared to MUT_ORF8. Energy minimized dimers obtained through MMGBSA were subjected to monomer interactions. From figure 3D and 3F it is clearly depicting that WT-ORF8 have 16 combined hydrogen bonds and salt bridges while MUT_ORF8 had only 8. Minimized dimers shows about difference of 2-fold in bond formation. These results clearly indicates that Mutant ORF 8 is losing its dimer formatting capacity which might affects the virus infectivity in the host.

**Figure 3:**
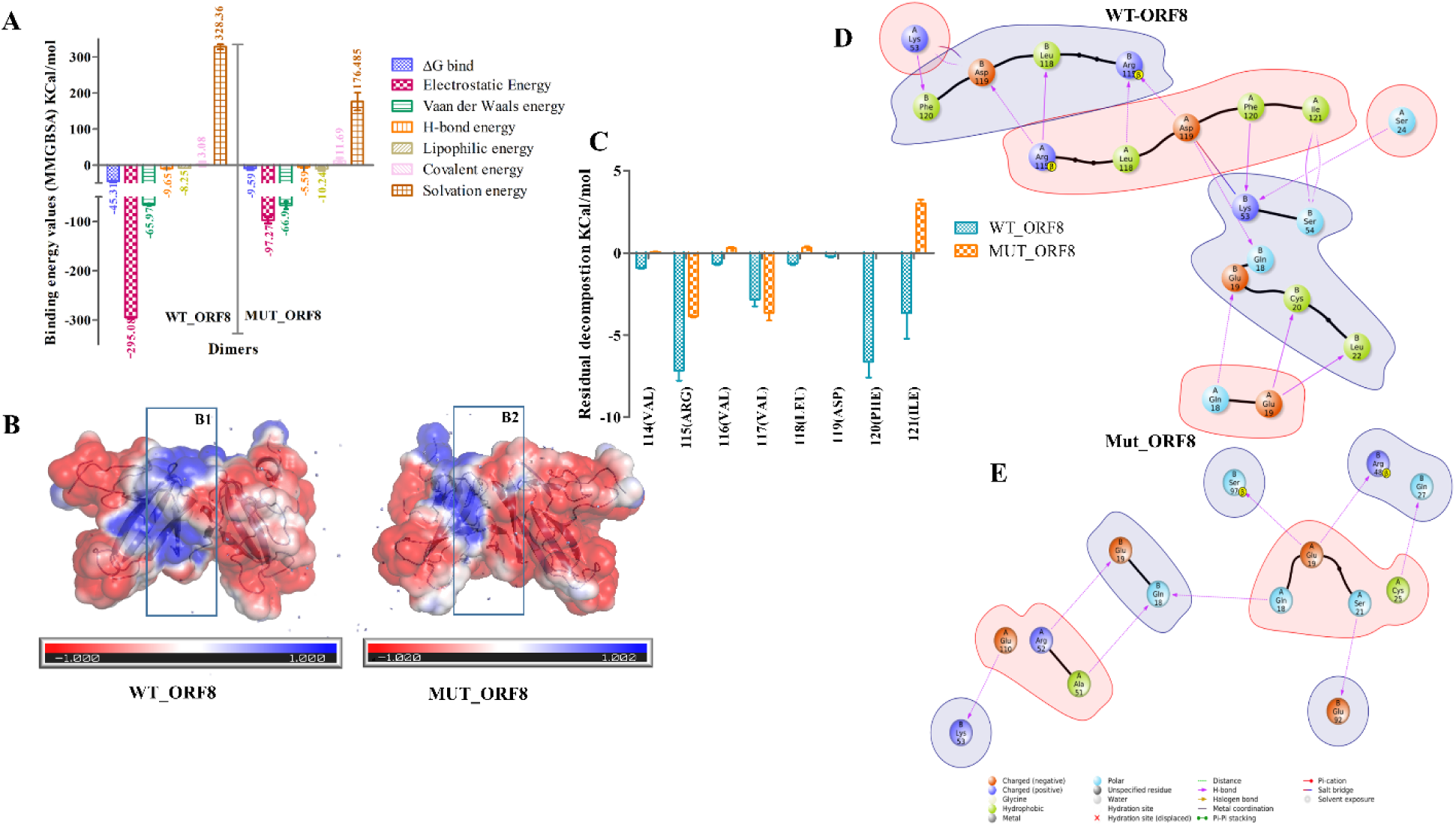
**B**inding energy studies within dimers of ORF8. **3A:** Binding energy difference between WT_ORF8 and MUT_ORF8. Major energies involved in dimer formation are shown in different legends. **3B:** electrostatic interaction map drawn for 1^st^ energy minimized dimer obtained from MMGBSA approach. Blue, white and red colors represent positive, null, negative electrostatic potential respectively, inform of surface representations. B1 & B2 represents potential between two monomeric subunits of WT-ORF8 and MUT-ORF8 respectively. **3C:** Thermal decomposition among amino-acids residues within both dimers. WT_ORF8 (cyan) and MUT_ORF8 (Orange) showing decomposition energies for key residues involved in dimer formations. **3D & 3E:** Interactions among energy minimized dimers obtained through MMGBSA, legends for each type of bond is shown in under figure 3E.

### 3.3 Flexible docking between Variants of ORF8 and MHC-I complex

As, the binding interface between ORF8 and MHC-I is not known yet, thus we used flexible docking to study the molecular interactions between ORF8 and MHC-I using PIPER. As shown in figure 4A, superimposed structure of docked pose of ORF8 and MHC-I were shown. Maximum possess which were generated were showing binding of ORF8 between beta macroglobulin chain and alpha 3 domain of MHC-I, where both dimers of ORF8 can easily accommodate. Pivotal interactions among WT_ORF8 with respect to MHC-I complex are 18 and MUT_ORF8 with respect to MHC-I were only 11 (Figure 4B & 4C). Based on docking results, we hypothesized that unstable dimeric structure of ORF8 (MUT_ORF8) might not be able to bind efficiently to MHC-I complex, hence not able to capture it tightly for autophagy. These correlations further lead to enhance expression of MHC-I compared to wild-type virus infection.

**Figure 4:**
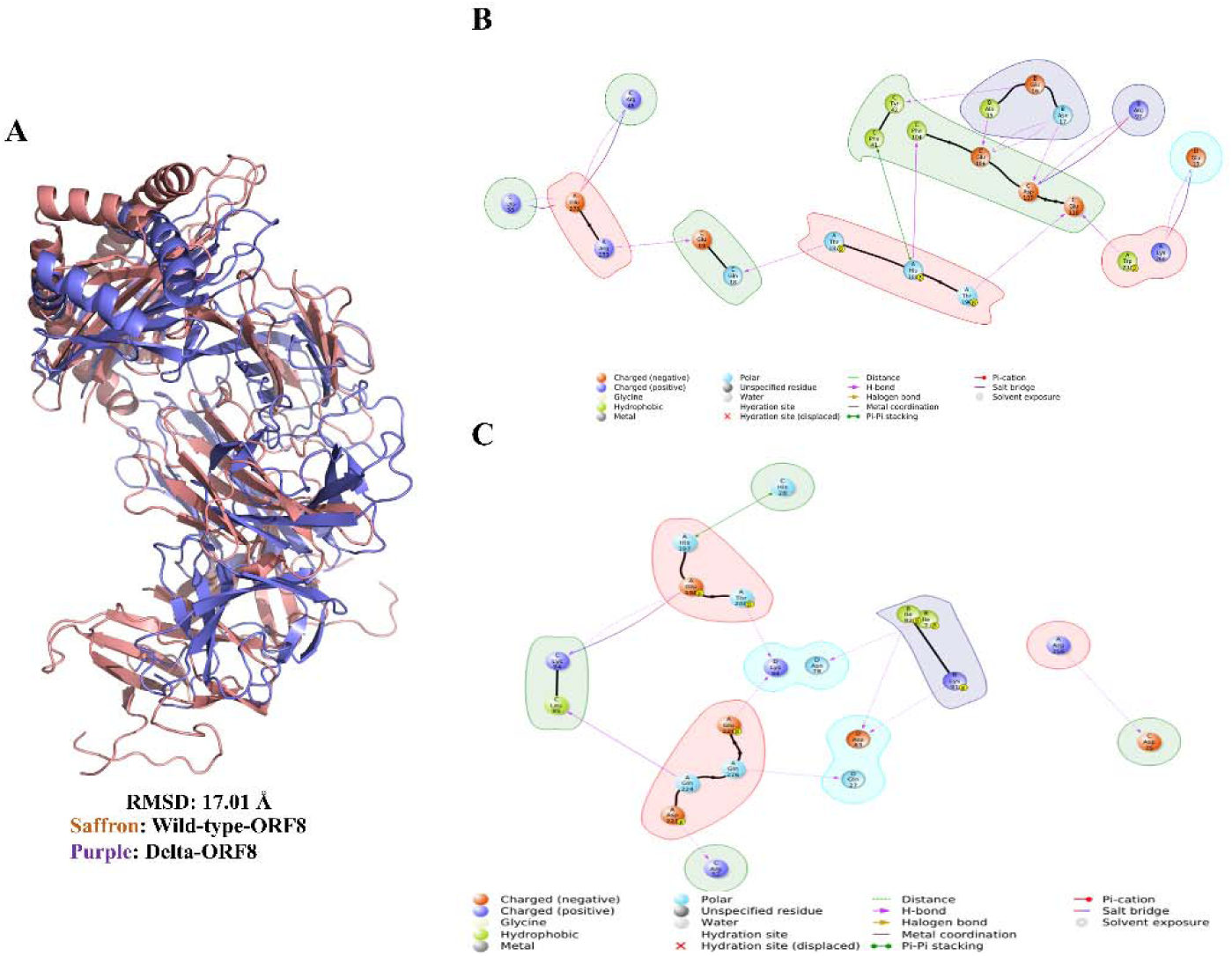
Flexible docking of MHC-I with ORF8 dimer. **A:** Superimposed structure of WT_ORF8_MHC-I(saffron) and MUT_ORF8_MHC-I (purple). **B:** Pivotal interaction among WT_ORF8_MHC-I complex. **C:** Pivotal interaction among MUT_ORF8_MHC-I complex.

## 4 Discussion

The molecular mechanism behind the severity and rapid spread of the COVID-19 disease is yet to be investigated. It is reported that ORF8 is a rapidly evolving dimeric protein that interfere with the immune responses in host [10]. There are some reports showing that ORF8 is interacting with proteins such as IL17RA of MHC-I molecular pathway [32]. It was also reported that SARS-CoV-2 virus infection leads to downregulation of MHC-I through direct interactions with ORF8 and selectively targeted towards lysosomal autophagy, consequently immune evasion [11]. The antigen presentation system of host will also be impaired due to ORF8-MHC-I interactions. So ORF8 has now become a prime target for scientist to investigate the mechanism behind ORF8-MHC-I interactions. During second wave of COVID-19 disease, although the infection rate was very high, it was seen that hosts developed adaptability towards the COVID-19 infection. Therefore, the study was planned with two objectives firstly, exhaustive analysis of the molecular structures of ORF8 dimer of wild type and delta variant (WT_ORF8 and MUT_ORF8) and secondly the interactions between WT_ORF8-MHC-I complex and MUT_ORF8-MHC-I complex. The detailed analysis of dimeric structures of WT_ORF8 and MUT_ORF8 showed a significant difference in interaction pattern between monomeric chain. In WT_ORF8 the key interaction is formed between Asp119 and Phe120 (Figure 1C). Whereas, due to deletion of Asp119 and Phe120 amino acids in MUT_ORF8 the interactions between MUT_ORF8 monomeric chains were diluted (Figure 1A). Deletion of Asp119 and Phe120 in MUT_ORF8 protein of SARS CoV2 delta variant caused loss of three salt bridges as well as H-bonds. The structural instability of the MUT_ORF8 can be clearly witnessed through molecular dynamics simulation studies. In MD studies RMSD, RMSF and radius of gyration of MUT_ORF8 is always towards higher side as compared to WT_ORF8 (Figure 2C, 2D, & 2F). It was also observed at many time points of simulation the number of hydrogen bonds tends to zero in MUT_ORF8 indicating that there was loss of connectivity between the monomeric chains of MUT_ORF8 (Figure 3D). But in WT_ORF8 there are constant interactions between the monomeric chains reveling the conformational stability of the dimeric structure. Higher RMSF values for MUT_ORF8 dimer throughout simulation indicates the greater flexibility. Additionally, the radius of gyration was also calculated for ORF8_WT and MUT_ORF8 dimers to study the compactness of these dimeric structure with protein folding and unfolding over thermodynamic principals during the 200ns of the molecular dynamics simulation. It is evident that only three amino acids i.e., In MUT_ORF8, amino acids Arg115, Ile119, Ala51, Ser24 are involved in bond formation between the dimers, whereas Phe120 and Lys53, Lys53 and Ser24, Gln18-Lue22, Arg52 and Ile121 in addition to A: Asp119-B: Arg115, A: ARG115-B: Glu92, B: Asp119-A: Arg115 and B: Arg115-A: Glu92 are involved in the stabilization of the WT_ORF8. Interestingly, in addition to these interaction two pi-Sulphur bonds were also observed between A: Phe120-B: Cys90 and A: Phe120-B: Cys25 in WT_ORF8, which is totally absent in MUT_ORF8 due to deletion of Phe120 amino acid. As Ala51 and Ser24 are major interacting amino acid in case of MUT_ORF8 its detail interaction map was built that surprisingly showed that it is these two amino acids are forming unfavorable bonds i.e. D: Ala51-D: Ser97 and C: Ser24-D: Lys53, which is also contributing towards instability of MUT_ORF8.

The stability of ORF8 dimers seems to be one of the major reasons contributing towards the host immune adaptability because the stable dimeric protein WT_ORF8 is able to tightly accommodate on the surface of MHC-I complex, whereas MUT_ORF8 is unable to firmly accommodate on the surface of MHC-I complex causing escape of MHC-I complex towards lysosomal autophagy and contributing consequently in increased immune response.

Nationwide population weighted study of seroprevalence from May-June 2020 was conducted by ICMR (Indian council of medical research), showing 0.75% among 21 states [33]. While in second seroprevalence study using Abbott assay detecting IGg antibodies against SARS-CoV-2 nucleoprotein, in August 2020 showed increased seroprevalence to 6.6% (95% CI 5·8–7·4) [34]. Seroprevalence among adults increased by about ten times, from 0·7% in May, 2020, to 7·1% in August, 2020 in India [35]. Supplementary figure 3A, showing number of SARS-CoV-2 cases reported in India during first and second wave. Third seroprevalence data shows percentage increase to 24.1% from December 2020 to January 2021 (Supplementary figure 3B). Drop down of 50% cases SARS-CoV-2 cases during second wave was double quick time compared to first wave. During first wave in 17^th^ September 2020 cases were 93735 and cases were halved in 6 weeks, 30^th^ October 2020 with 46380 cases (Supplementary figure 3C). While in second wave higher number of cases (3, 91,261) where decreased (1, 95,183) in half time compared to first wave. ICMR 4^th^ seroprevalence data shows 70% of Indian population (unvaccinated) showing IgG antibody titer against SARS-CoV-2 cases (ICMR 5^th^ Seroprevalece data). Drastic increase in seroprevalence after second wave, from 0.75% to 70% is unusual observation for high transmittable delta variant. Delta variant have these D119, F120 deletions, which were disrupting ORF8, responsible for downregulating MHC-I and suppressing host immune response. Results of the study suggests that the dimerization of MUT_ORF8 is altered, that might be affecting the ORF8 mediated MHC-I downregulation by autophagy in delta variant.

Second in India was not only due to predominant delta variant but other lineages were also involved. In such cases strong case study or proof is required in support of this hypothesis that antibody response was due to loss of dimerization capacity of ORF8. In nationwide study seroprevalence was detected for all kind of SARS-CoV-2 lineage infections, but here we are studying only delta mediated immune responses. To support this nationwide study, State wise seroprevalence was also studied in region like Ahmedabad, Gujarat having higher number of active cases of SARS-CoV-2 infection. Supplementary figure shows the genome sequencing data of Gujarat biotechnology research center during second wave, where B.1.167.2 (red) lineage (delta) was found to be 100% in samples collected from patients [37]. 5^th^ seroprevalence data of Ahmedabad city is shown in Supplementary figure S4B. Seroprevalence due to delta only was 81.93% (Ahemdabad Summary, 2021). We had hypothesized that altered dimer of ORF8 might not able to perform autophagy of MHC-I molecule compared to wild-type ORF8, which might lead to favoring host immune responses. This can be one possible reason for the sudden drop down of cases during second wave in India.

## 5 Conclusion

Frequency of delta variant during second wave in India was persisted 9.6-76.5% in India while in Gujarat it was between 18.96 to 90% (Figure 5D). 5^th^ seroprevalence study by ICMR shows 62.3% population have antibodies due to virus infection, while in Gujarat there 81.93% seroprevalence was observed. These patterns leads to conclude that as the frequency of delta is increasing seroprevalence among population had also increased (Figure 5C & 5D). These seroprevalence study supports our hypothesis that loss in dimerization capacity of ORF8 (from delta variant) leads to an abrogation of ORF8 MHC-I interaction and overcome suppression of adaptive immune response.

**Figure 5:**
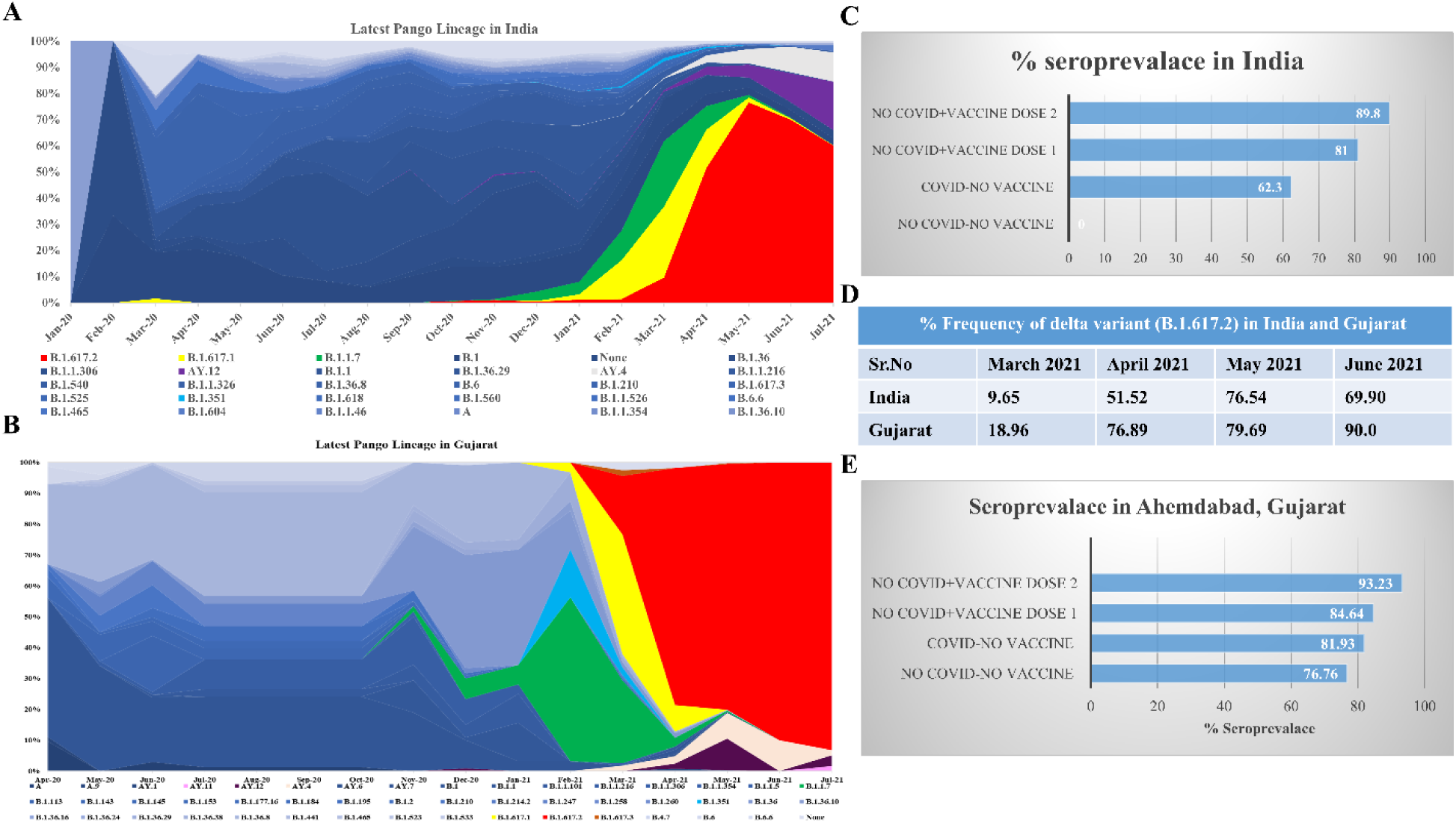
Nationwide and statewide seroprevalence study: **5A:** SARS-CoV-2 sequences submitted to GAISAD database from India at different time scale with latest Pango lineage. **5B:** SARS-CoV-2 sequences submitted to GAISAD database from Gujarat at different time scale with latest Pango lineage. **5C:** 5^th^ Seroprevalence data from ICMR (Indian council of medical research). **5D:** Table narrating frequency of delta variant (B.1.617.2) during second wave in India and Gujarat. **5E:** 5^th^ Seroprevalence data FROM Ahmedabad city, Gujarat.

## 7. Authors contributions

AC and IS performed Insilco experiments, Molecular dynamics, validated hypothesis and wrote the manuscript. AP, MJ, and CJ provided funding, validated results, and corrected manuscript.

## 8. Data Availability Statement

Seroprevalence study were performed based on ICMR (Indian council of medical research), PIB (Press information bureau), Times of India (TOI), Gujarat Biotechnology research center’s COVID19 portal (GBRC), MoHFW (Ministry of health and family welfare), Nextstrain and GISAID data base.

ICMR: https://www.icmr.gov.in/

PIB: https://pib.gov.in/PressReleseDetail.aspx?PRID=1748351

TOI: https://timesofindia.indiatimes.com/

GBRC: https://covid.gbrc.res.in/

MoHFW: https://www.mohfw.gov.in/

Nextstrain: https://nextstrain.org/ncov/gisaid/global

GISAID: https://www.gisaid.org/index.php?id=209

