## Supplemental figures and video for "Defective ORF8 dimerization in delta variant of SARS CoV2 leads to abrogation of ORF8 MHC-I interaction and overcome suppression of adaptive immune response": Supplementary paper for figures_FROORF8.docx

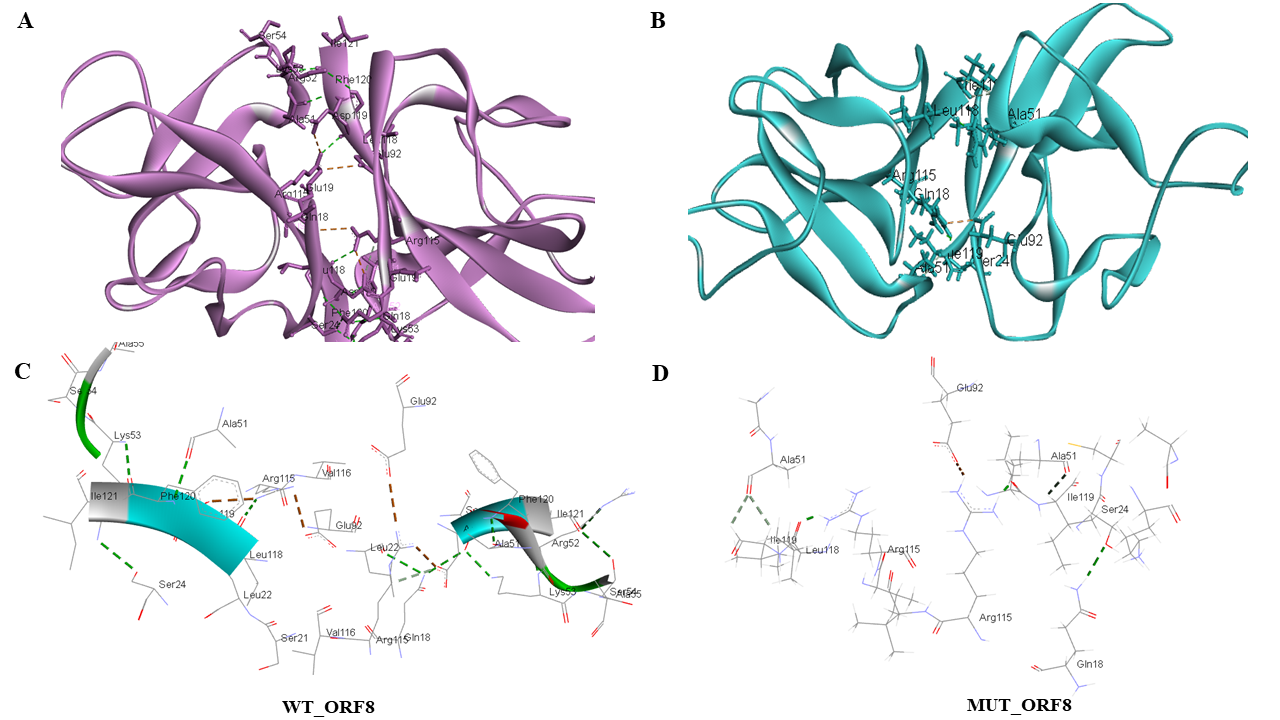

**Supplementary figure S1:** Wild-type (WT_ORF8) and Mutant (MUT_ORF8) ORF8 interactions showing salt bridges (orange) and hydrogen bonds (red) within 3.5 distance visualized in Biovia discovery studio. **A & B:** WT_ORF8 (Magenta) and MUT_ORF8 (cyan) cartoon representation showing salt bridges and hydrogen bonds, respectively. **C & D:** WT_ORF8 stick representation showing salt bridges and hydrogen bonds, where one can visualize higher bonding in WT compare to mutant respectively.

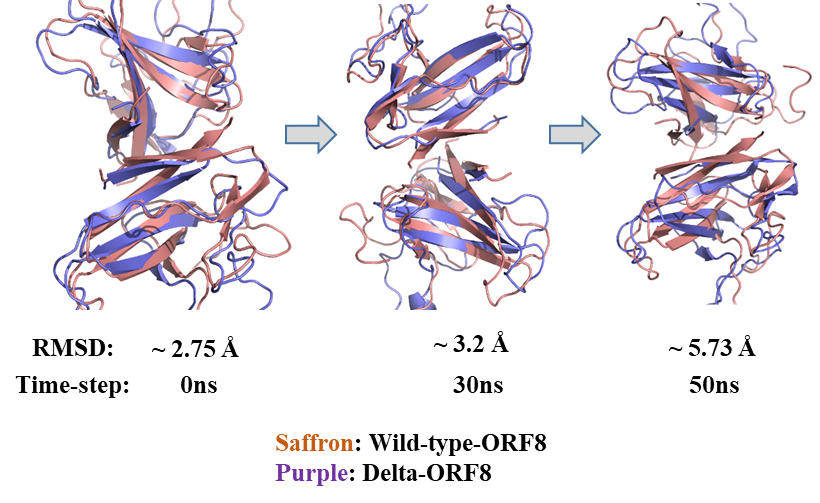

**Supplementary figure S2:** Important protein conformational dynamics during various time point.

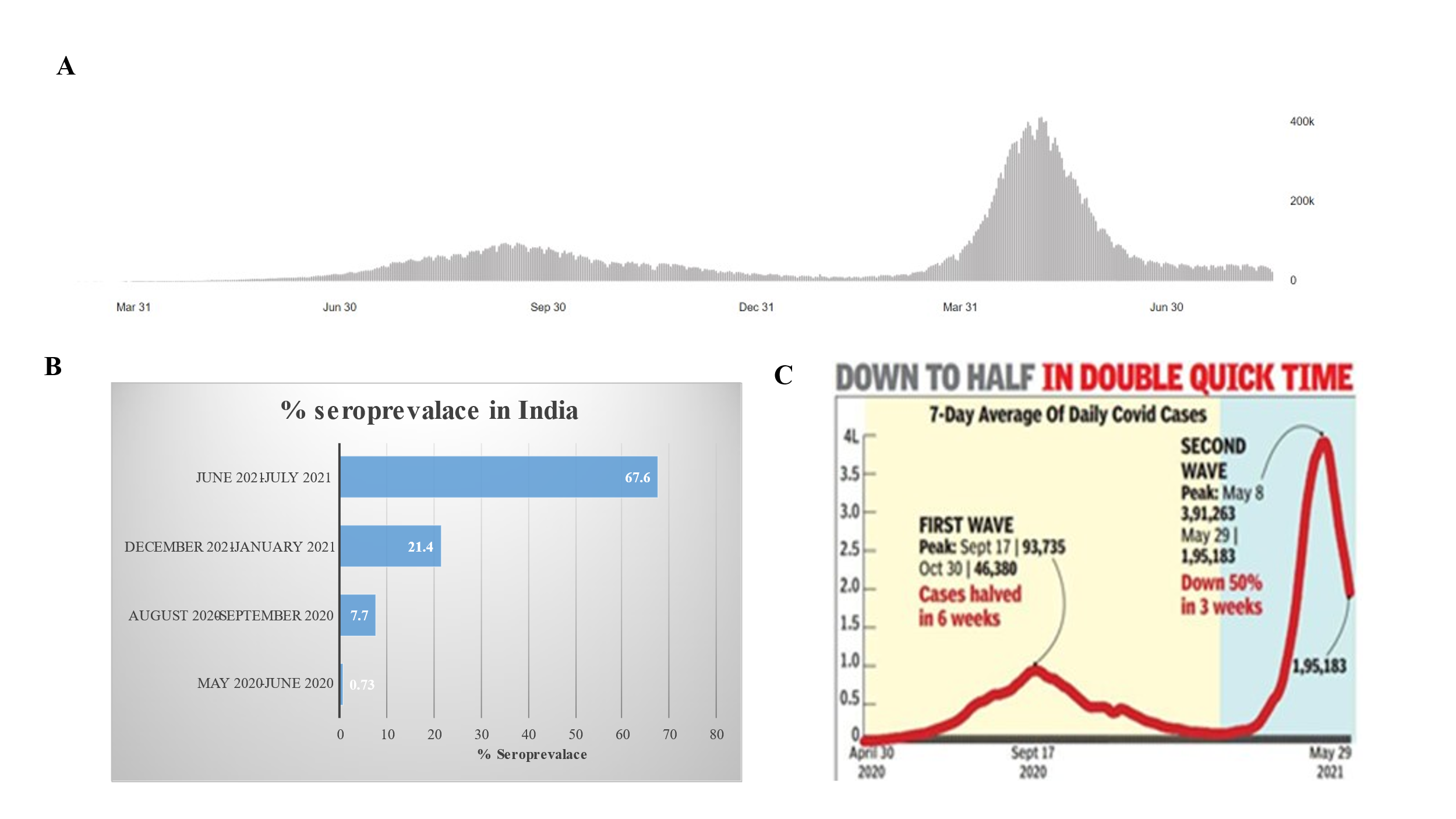

**Supplementary figure S3:** Seroprevalence study during SARS-CoV-2 pandemic in India. **A:** Number of cases reported during SARS-CoV-2 infecting during first and second wave. **B:** % Seroprevalence reports studied by ICMR. **C:** Comparison of Cases pattern during SARS-CoV-2 infection in India report by TOI (Times of India)

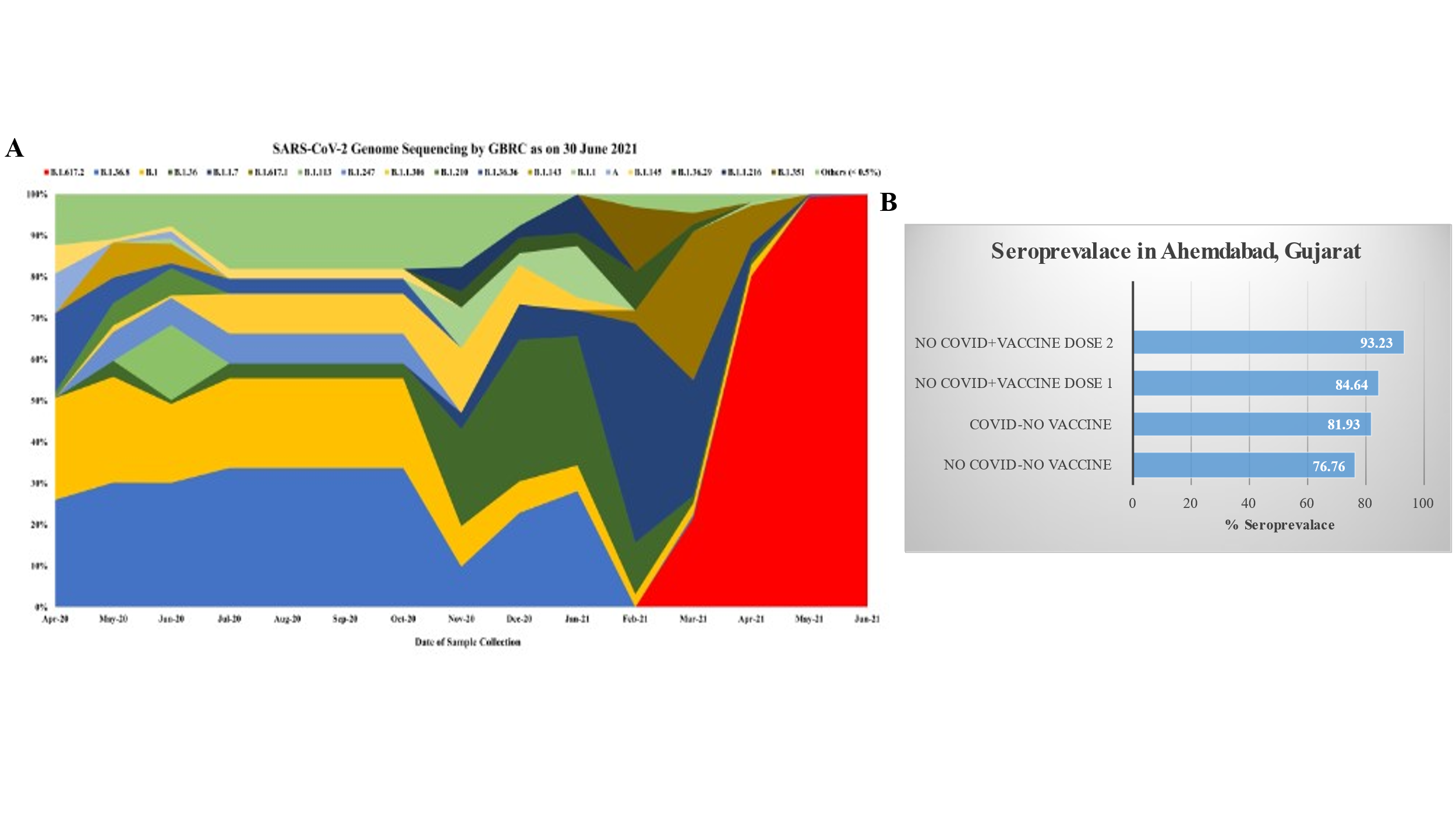

**Supplementary figure S4:** seroprevalences in Gujarat. **A.** Genome sequencing done by Gujarat Biotechnology Research Center showing SARS-CoV-2 variants during second wave. **B.** 5^th^ seroprevalence data of Ahmedabad city during second wave.

**Supplementary Video 1:** Wildtype ORF8

**Supplementary video 2:** Mutant ORF8
